# LncRNA-TUG1/EZH2 axis promotes cell proliferation, migration and the EMT phenotype formation through sponging miR-382

**DOI:** 10.1101/123141

**Authors:** Liang Zhao, Hongwei Sun, Hongru Kong, Zongjing Chen, Bicheng Chen, Mengtao Zhou

## Abstract

Pancreatic carcinoma (PC) is the one of the most common and malignant cancer in the world. Despite many effort have been made in recent years, the survival rate of PC still remains unsatisfied. Therefore, investigating the mechanisms underlying the progression of PC might facilitate the development of novel treatments that improve patient prognosis. LncRNA Taurine Up–regulated Gene 1 (TUG1) was initially identified as a transcript up - regulated by taurine, siRNA - based depletion of TUG1 suppresses mouse retinal development, and the abnormal expression of TUG1 has been reported in many cancers. However, the biological role and molecular mechanism of TUG1 in pancreatic carcinoma (PC) still needs to be further investigated. In the current study, the expression of TUG1 in the PC cell lines and tissues was measured by quantitative real-time PCR (qRT-PCR), and loss-of-function and gain-of-function approaches were applied to investigate the function of TUG1 in PC cell. Online database analysis tools showed that miR-382 could interact with TUG1 and we found an inverse correlation between TUG1 and miR-382 in PC specimens. Moreover, dual luciferase reporter assay, RNA-binding protein immunoprecipitation (RIP) and applied biotin-avidin pulldown system further provide evidence that TUG1 directly targeted miR-382 by binding with microRNA binding site harboring in the TUG1 sequence. Furthermore, gene expression array analysis using clinical samples and RT-qPCR proposed that EZH2 was a target of miR-382 in PC. Collectively, these findings revealed that TUG1 functions as an oncogenic lncRNA that promotes tumor progression at least partially through function as an endogenous ‘sponge’ by competing for miR-382 binding to regulate the miRNA target EZH2.

## Introduction

Pancreatic cancer (PC) is one of the deadliest human malignancies and the fourth-leading cause of cancer-related deaths[1, 2]. Due to lacking efficient methods for early diagnosis, patients are always diagnosed as advanced stage. Currently, surgical resection is important treatment for PC; however, only 10-20% patients are diagnosed at a resectable stage and the 5-year survival of post-operation for PC is still unsatisfied with only 20%[3]. Thus, chemotherapy becomes one of the indispensable means of clinical treatment of PC. Early metastasize is one of the characteristics of pancreatic cancer, which always result in poor curative effect or failure of chemotherapy. It has been reported that epithelial-to-mesenchymal (EMT), process of losing epithelial phenotypes and gaining of mesenchymal phenotypes, contributes to the early dissemination of primary tumor cells and metastatic spread[4-6]. The aim of our present study is to explore the underlying EMT formation mechanism of PC and potential therapeutic value.

Long non-coding RNA (lncRNA)s, defined as non-coding RNA more thatn 200nt with limit of no coding protein ability, are gaining prominence. The function of lncRNAs has been characterized in many cancers. LncRNAs exert their function mainly at three levels: transcription, post-transcription and epigenetic modification. Taurine-upregulated gene 1 (TUG1), initially identified as a transcript that upregulated in response to taurine treatment of developing mouse retinal cells, has been reported aberrantly expressed in many human cancers and involved in carcinogenesis and progression[7-10].

In our present study, we explored the role of TUG1 in the regulation of proliferation, migration and EMT formation of PC. Our findings show that silencing of TUG1 impairs migration and epithelial mesenchymal transition ability. Moreover, for the first time, we identify an important role for TUG1 in pancreatic cancer cells. Furthermore, we demonstrate that TUG1could competitive sponging miR-382 and then regulating the expression of EZH2, therefore involving in the progression of PC.

## Methods and materials

### Patients

Pancreatic samples were obtained from the Department of Surgery, The First Affiliated Hospital, Wenzhou Medical University. All patients did not receive chemotherapy or radiotherapy before the surgery. All samples were frozen in liquid nitrogen immediately after resection and stored at −80 °C until use. This study was approved by the institutional review boards of all of the hospitals involved in the study, and the written informed consent was obtained from all patients.

### Cell lines

Human PDAC cell lines PANC-1, AsPC-1 and HEK 293T cell were obtained from ATCC (the American Type Culture Collection, Manassas, VA, USA). Patu 8988, BxPC-3, SW 1990 cells and normal pancreatic cells HPDE6-C7 were purchased from the Institute of Biochemistry and Cell Biology of the Chinese Academy of Sciences(Shanghai, China). All cell lines were cultured in 37 °C in a 5% CO2 humidified in DMEM or RPMI 1640 (Gibco) with 10% fetal bovine serum (FBS, HyClone).

### Cell cytoplasm/nucleus fraction isolation

NE-PER Nuclear and Cytoplasmic Extraction Reagents (Thermo Scientific, Waltham, MA, USA) was employed to prepare cytoplasmic and nuclear extracts from PDAC cells. RNAs extracted from each of the fractions were subjected to following RT–qPCR analysis to demonstrate the levels of nuclear control transcript (MALAT1), cytoplasmic control transcript (GAPDH), lncRNA TUG1, and miR-382.

### Quantitative real time PCR Analyses

The total RNA was extracted from the tissues or cultured cells using TRIzol reagent (Invitrogen, Carlsbad, CA, USA), according to the manufacturer’s instructions. Reverse transcription was performed with PrimeScript RT reagent Kit (Takara, Dalian, Liaoning, China). qRT-PCR was performed with SYBR Prime Script RT-PCR Kits (Takara, Japan) based on the manufacturer’s instructions. The target gene expression level was calculated with the 2^-ΔΔct^ method, which was normalized to GAPDH mRNA. All assays were performed in triplicate. The expression levels were relative to the fold change of the corresponding controls, which were defined as 1.0.

### Cell viability

Cell viability was assessed via 3-(4,5-dimethylthiazol-2-yl)-2, 5-diphenyl-trtrazolium bromide (MTT) assay. 5 × 103 cells/well were seeded in a 96-well flat-bottomed plate for 24 h, then transfected with corresponding sh-RNA or pc-DNA3.1 and cultured in normal medium. At 0, 24, 48, 72 h and 96h after transfection, the MTT solution (5 mg/ml, 20 μl) was added to each well. Following incubation for 4 h, the media was removed and 100 μl DMSO were added to each well. The relative number of surviving cells was assessed by measuring the optical density (O.D.) of cell lysates at 560 nm. All assays were performed in triplicate.

### Colony formation assay

The indicated cells were plated into 6-well plates (1*10^3^ cells/well) and incubated in DMEM or RPMI 1640 with 10% FBS at 37 °C. Two weeks later, cells were washed with PBS, fixed in methanol for 30 min and stained with 1% crystal violet dye, and the number of colonies was counted. All assays were performed in triplicate.

### Cell migration and invasion assays

Cell migration were measured by transwell chamber (8 um pore size, Corning). 48 h after transfection, cells in serum-free media were placed into the upper chamber. Media containing 10% FBS was added into the lower chamber. Following 48 h incubation, cells remaining in upper membrane were wiped off, while cells that migrated were fixed in methanol, stained with 0.1% crystal violet and counted under a microscope. Three independent experiments were carried out.

### Wound healing assays

Cell migration capacity was calculated by wound healing assay. 2× 10^5^ cells with or without transfection were plated into 12-well plates and incubated in DMEM or RPMI 1640 with 10% FBS at 37°C. After reaching 100% confluence, cells were wounded by scraping with a 200µl tip, following washed 3 times in serum-free medium and incubated in regular medium. Wounds were observed at 0 and 48 h. The cell migration distance was calculated by subtracting the wound width at each time point from the wound width at the 0h time point. Three independent assays were assayed.

### Western blots

Cells were lysed in RIPA lysis buffer (P0013, Beyotime) and nuclear proteins were extracted using lysis buffer (P0028, Beyotime), all the procedures were following the manufacturer’s protocol. Subsequently the cell lysates were boiled in 5X SDS-PAGE loading buffer for 10 min and then resolved by 8% SDS-PAGE and transferred to nitrocellulose membrane. The following antibodies were used in this study: E-cadherin, N-cadherin, Vimentin, EZH2 and β-catenin were purchased from Cell signaling technology. GAPDH (Proteintech) was referred as internal reference. Bound antibodies were visualized with the ECL kit (P0018, Beyotime).

### Dual luciferase reporter assay

Cells (293 T) were seeded at 3 × 10^4^ cells/well in 24-well plates and allowed to settle overnight. The next day, cells were co-transfected with pmirGLO-TUG1-WT or -MUT reporter plasmids and miR-382 mimic. And also cells were co-transfected with pmirGLO-EZH2-WT or -MUT reporter plasmids and miR-382 mimic. Twenty-four hours after transfection, the relative luciferase activity was measured using the Dual-Luciferase Reporter Assay System (Promega, Madison, WI, USA) and normalized against Renilla luciferase activity.

### RNA-binding protein immunoprecipitation

RIP experiments were performed using the Magna RIP RNA-binding protein immunoprecipitation kit (Millipore, Billerica, MA, USA) and the Ago2 antibody (Abcam, Cambridge, MA, USA) following the manufacturer’s protocol. Co-precipitated RNAs were subjected to RT–qPCR analysis.

### RNA-Pulldown assay

PANC-1 cells were transfected with biotinylated miRNA, collected 48 h after transfection. The cell lysates were incubated with M-280 streptaviden magnetic beads (Invitrogen, San Diego, CA, USA). The bound RNAs were purified using TRIzol reagent (Invitrogen) for further RT–qPCR analysis.

### Statistical analysis

Statistical analyses were performed using SPSS 17.0 software (SPSS, Chicago, IL, USA). Differences between two groups were assessed using Student’s t-test (two-tailed). Correlations between TUG1 and miR-382 or EZH2 were analyzed by Spearman rank correlation. Each experiment was performed at least three times. Results of experiments are displayed as mean ± SD. A P-value<0.05 was considered to indicate statistical significance.

## Results

### TUG1 is upregulated in PC and correlated with clinical outcome of pancreatic cancer patients

To study the lncRNAs mechanism involved in regulating the progression of PC, we previously conducted lncRNA expression array analysis using clinical samples (Fig. 1A). Among those lncRNAs, lncRNA-PVT1 and lncRNA-TUG1 were highly expressed in pancreatic tissues. The biology function of PVT1 has been discussed in our other study. In the present study, we focus on the function of TUG1. To further validate the expression level of TUG1, we evaluated the TUG1 expression level in 34 paired PC and adjacent pancreatic tissue samples using reverse transcription and quantitative real-time PCR (RT–qPCR). The results showed that TUG1 levels were significantly higher in tumor tissues (Fig. 1B). Additionally, correlation between lnc-TUG1 expression and clinical features revealed high level of TUG1 was correlated with large tumor size (p=0.000), poor tumor differentiation (p=0.000), TNM stage (p=0.001), Vascular infiltration (p=0.000), distant metastasis (p=0.000) and negatively correlated with overall survival (OS) of patients with pancreatic cancer (Figure 1C), which indicated that upregulated TUG1 might contribute to the development of pancreatic cancer. Then, we examined the expression level of TUG1 in PC cell lines (PATU 8988, BxPC-3, PANC-1, SW 1990, AsPC-1) and HPDE6-C7 cells, an immortalized, normal human pancreatic normal pancreatic epithelial cell line, using RT–qPCR. Compared with HPDE6-C7 cells, PC cells exhibited significantly higher levels of TUG1 expression but not in PATU 8988 cells (Figure 1D). Collectively, these results suggest that TUG1 is up-regulated in PC.

**Figure 1.**
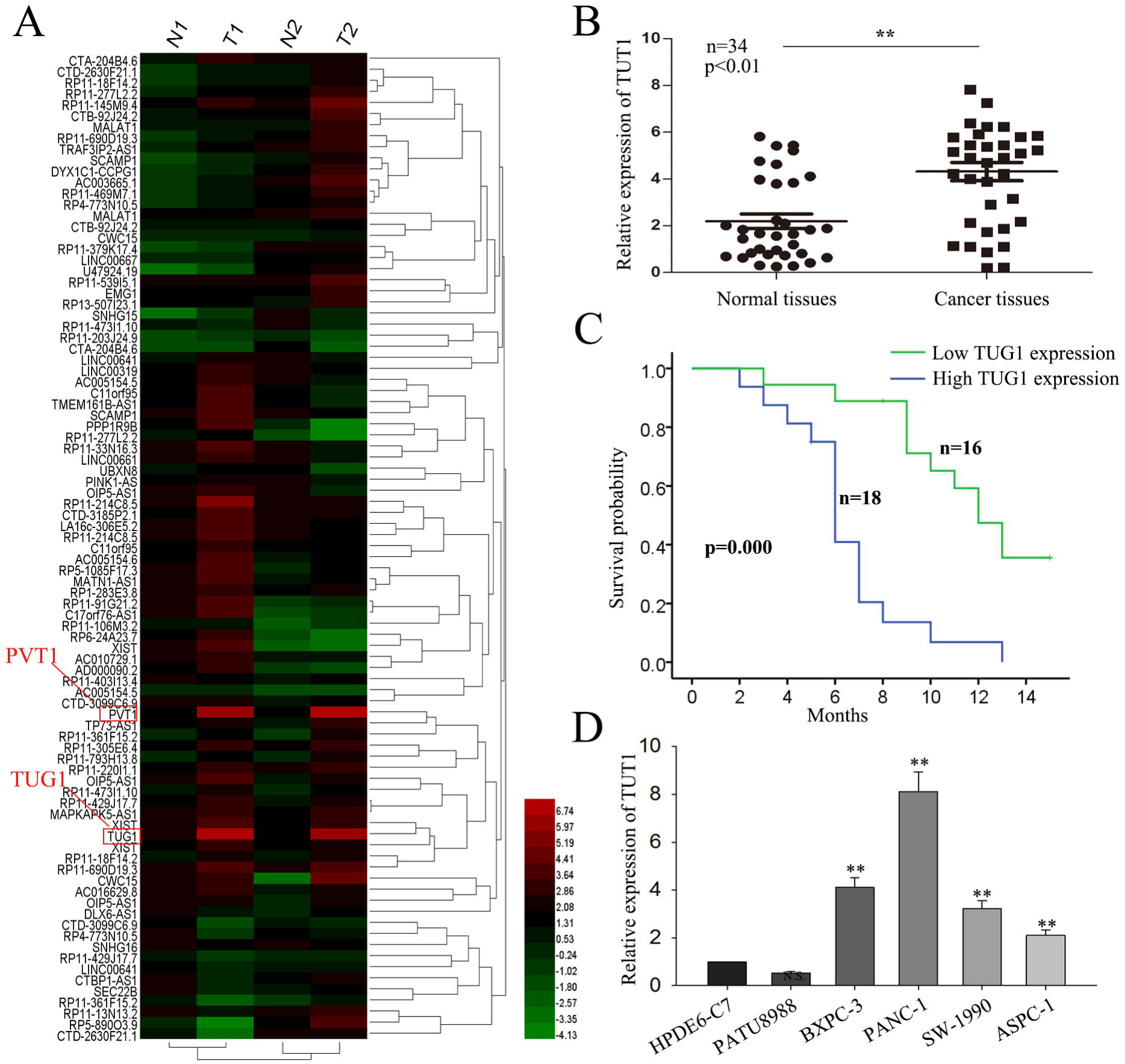
TUG1 is upregulated in PC and correlated with clinical outcome of pancreatic cancer patients. **(A)** LncRNA expression array analysis of clinical samples. **(B)** TUG1 expression in pair samples of PC and adjacent normal tissues. **(C)** The overall survivals in 34 pancreatic cancer (PC) patients were represented by Kaplan-Meier curves. **(D)** Relative levels of TUG1 in normal HPDE6-C7 cells and five types of PC cells. The p-value represents the comparison between groups (*p < 0.05, **p < 0.01).

### TUG1 promotes PC cell proliferation, migration and EMT formation

To evaluate the role of TUG1 in regulating the proliferation, migration and EMT formation, we transfected PATU 8988 and PANC-1 cell lines with TUG1expression vector and shRNA/TUG1, respectively (Fig. 2A). First we performed the MTT and colony formation assay to monitor the effect of TUG1 on cell proliferation. We observed a significant increase in proliferation of cells transfected with TUG1expression vector compared with negative control in PATU 8988 cells (Fig. 2B and 2C, left). Furthermore, down-expression of TUG1 in PANC-1 cells resulted in a significant increase in cell proliferation (Fig. 2B and 2C, right). Then, we test the effect of TUG1 on cell migration capacity. Wound healing assay showed that forced expression of TUG1 drastically increased the cells migration capacity, while TUG1 knockdown significantly decreased the cells migration capacity (Fig. 2D-E). These data provide evidence of the migration-promoting role of TUG1 in vitro. Additionally, we examined the influence of TUG1 on the formation of EMT. Results from western blot showed that overexpression of TUG1 contributed to the formation of EMT, and vice versa in cells deletion of TUG1 (Fig. 2F-G). Taken together, these results suggest that TUG1 has important roles in PC cells proliferation, migration and EMT formation. However, the detailed mechanism by which TUG1 functions warrants further investigations.

**Figure 2.**
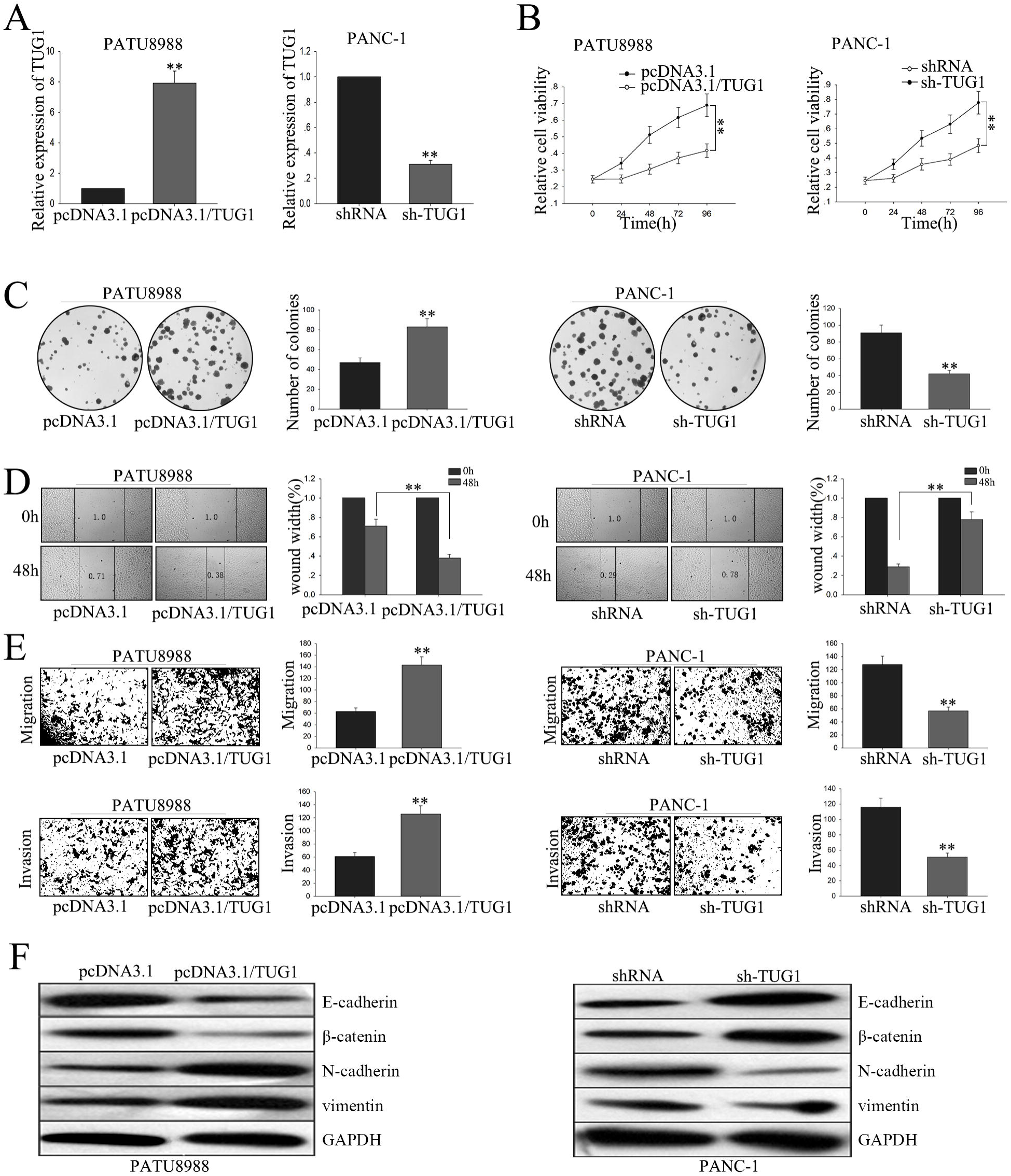
TUG1 promotes PC cell proliferation, migration and EMT formation. **(A)**The overexpression and downexpression efficiency of TUG1 expression vector and sh-TUG1 was obtained after transfection 48h. **(B)** The MTT, **(C)** colony formation, **(D)** wound healing assay, **(E)** transwell assay, **(F)** western blot assay was performed in PATU 8988 cells transfected with pcDNA3.1/TUG1or in PANC-1 cells with sh-TUG1. All data were represented as the mean ± S.D. from three independent experiments, *P < 0.05, **P < 0.01.

### MiR-382 is regulated by TUG1

Besides characterizing the function of TUG1 on PC cells, we also sought to explore in further detail the mechanism by which TUG1 exerts its functions. Accumulating documents identified that many lncRNAs have been reported to function as competing endogenous (ce)RNAs by serving as sponges that bind and sequester away miRNAs[11-14]. To exam whether TUG1 exerts its function in PC cells through function as a ceRNA, we first measured the percentage of TUG1 in the cytoplasmic and nuclear fractions of PANC-1 cells. As shown in Fig. 3A, RT–qPCR of nuclear and cytoplasmic fractions of PANC-1 cells present that TUG1 was located in the cytoplasm, providing prerequisite for function as a ceRNA. Then, using starbase software (http://starbase.sysu.edu.cn/mirLncRNA.php), we measured the miRNAs predicted to be bound to TUG1 in response to knockdown of TUG1. We transfected PANC-1 cell with shTUG1, and we found that down-regulated TUG1 had the most significant effect on miR-382 expression (Supplementary Table1). Next, RT–qPCR of nuclear and cytoplasmic fractions of PATU 8988 and PANC-1 cells validated that miR-382 was also located in the cytoplasm, indicating a potential reciprocal interaction between TUG1 and miR-382 (Fig. 3B). Thus, miR-382 was pursued for studies in detail. RT–qPCR showed that, in PC tissues, miR-382 was downregulated in the tumor specimens, which was inversely correlation with TUG1 (Fig. 3C). Compared with HPDE6-C7 cells, PC cell lines displayed lower expression of miR-382, except for in PATU 8988 cells (Fig. 3D). To further evaluate the potential regulating relationship between TUG1 and miR-382, PATU 8988 and PANC-1 cells were transfected with TUG1expression vector and shRNA/TUG1 to increase or decrease TUG1 expression, respectively. Forced expression of TUG1 resulted in a significant down-regulation of miR-382, while deletion of TUG1 significantly increased the level of miR-382 (Fig. 3E). These results suggest that miR-382 was negatively regulated by TUG1. To further explore whether TUG1-mediated miR-382 regulation through function as a ceRNA, we performed dual luciferase reporter assay, RNA-binding protein immunoprecipitation (RIP) and applied biotin-avidin pulldown system. We first sub-cloned full-length of TUG1 into the pmirGLO dual luciferase reporter vector and performed luciferase assays in PANC-1 cells (Fig. 3F). As shown in Fig. 3F, co-transfection of PANC-1 cells with pmirGLO-TUG1-WT vector and miR-382 mimic significantly reduced luciferase reporter activity with respect to the negative control, while cells co-transfected with pmirGLO-TUG1-MUT vector and negative control showed a higher level of luciferase activity in comparison to the group of co-transfected with WT vector and negative control. Results from RIP showed that Ago2 protein was efficiently immunoprecipitated from cell extracts by Ago2 antibody (Fig. 3G), which revealed that while TUG1was detected in Ago2 immunoprecipitates from the control group, its levels were drastically reduced in Ago2 complexes purified from cells treated with miR-382 inhibitor, indicating that indicating that TUG1 is likely in the miR-382 RISC complex. Further, a biotin-avidin pulldown system was application to assay whether miR-382 could pull down TUG1. As shown in Fig. 3H, TUG1 was pulled down by miR-382, but the bind site of TUG1 for miR-382 was mutated failure to pull down TUG1, indicating that TUG1 regulated miR-382 in a sequence-specific manner. Together, our findings revealed that TUG1 exerts inhibitory effects on miR-382 expression through function as a ceRNA and therefore directly sponging miR-382.

**Figure 3.**
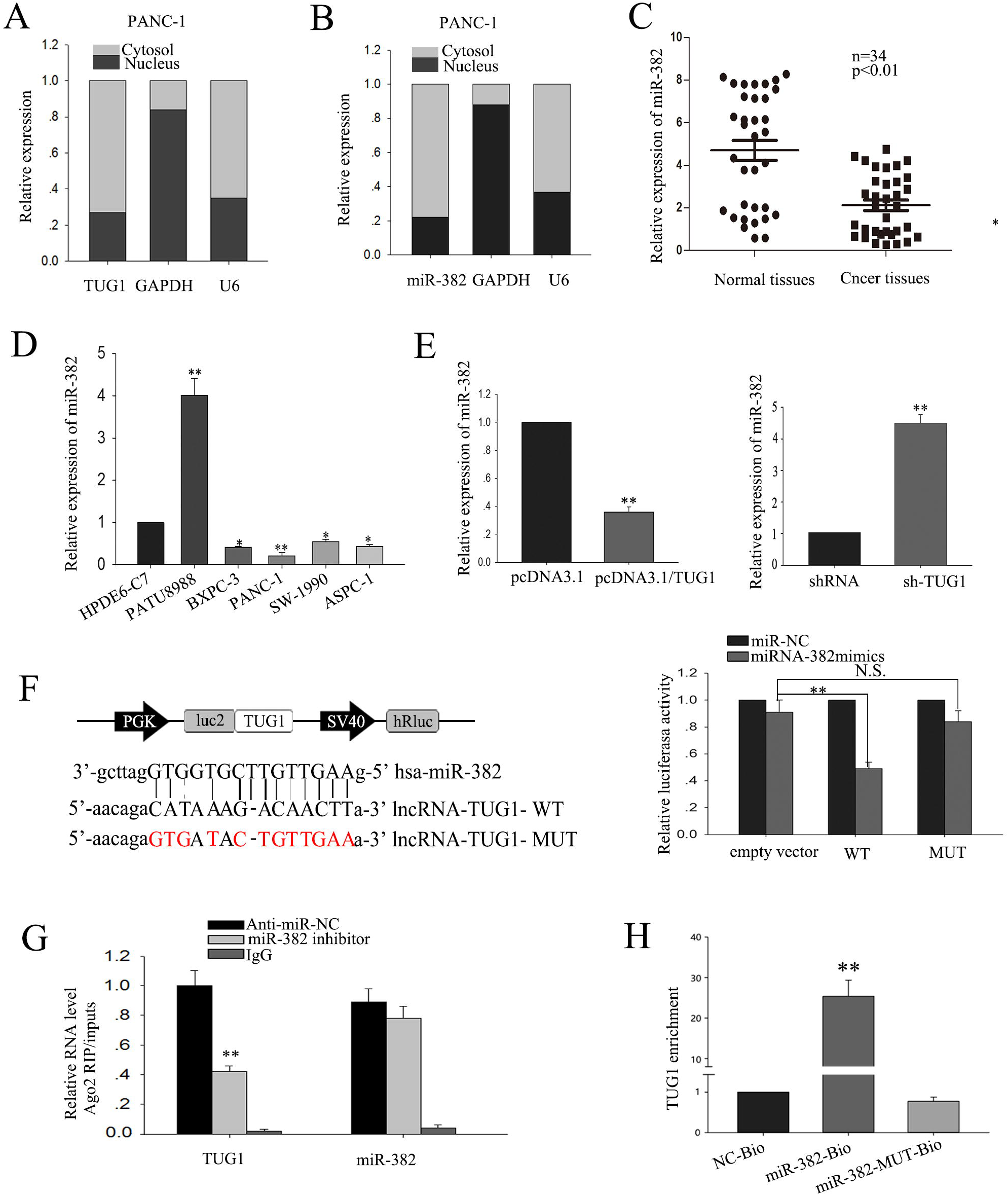
MiR-382 is regulated by TUG1. (**A)** RT–qPCR detection of the percentage of TUG1, miR-382 **(B)**, GAPDH and U6 in the cytoplasmic and nuclear fractions of PANC-1 cells. GAPDH and U6 serve as a cytoplasmic and nuclear localization marker, respectively. **(C)** miR-382 expression in pair samples of PC and adjacent normal tissues. **(D)** Relative levels of miR-382 in normal HPDE6-C7 cells and five types of PC cells. **(E)** The expression of miR-382 in response to the level of TUG1. **(F)** Luciferase assays of the indicated cells transfected with pmirGLO-TUG1-WT or pmirGLO-TUG1-MUT reporter and NC or miR-382 mimic. **(G)** Amount of TUG1 bound to Ago2 or IgG measured by RT–qPCR after RIP, IgG was used as a negative control and SNRNP70 was used as a positive control. **(H)**. PANC-1 cells were transfected with biotinylated WT miR-382 (miR-382-Bio) or biotinylated mutant miR-382 (miR-382-MutBio) or biotinylated NC (NC-Bio). Forty-eight hours after transfection, cells were collected for biotin-based pulldown assay. TUG1 expression levels were analyzed by RT–qPCR. All data were represented as the mean ± S.D. from three independent experiments, *P < 0.05, **P < 0.01.

### MiR-382 reversed the function of TUG1 in PC cells

We transfected PATU 8988 with TUG1expression vector and miR-382 mimic to study the effects of miR-382 on cell proliferation, migration and EMT mediated by TUG1. MTT and colony formation assay revealed that TUG1 promoted and miR-382 suppressed cell proliferation, while co-transfection of miR-382 mimic and TUG1 expression vector showed that TUG1 promoted cell proliferation was inhibited by miR-382 (Fig. 4A-B). Would healing assay and transwell assay revealed that TUG1 promoted and miR-382 suppressed cell migration, while co-transfection of miR-382 mimic and TUG1 expression vector showed that TUG1 promoted cell migration was inhibited by miR-382 (Fig. 4C-D). Moreover, results from western blot demonstrated that TUG1 promoted and miR-382 suppressed PC cell EMT phenotype formation, while co-transfection of miR-382 mimic and TUG1 expression vector showed that miR-382 reversed the EMT promoted by TUG1 to MET (Fig. 4E). Those observations suggest that the effects of TUG1 overexpression on the promotion of PC cell migration and EMT formation could be reversed by miR-382 mimics.

**Figure 4.**
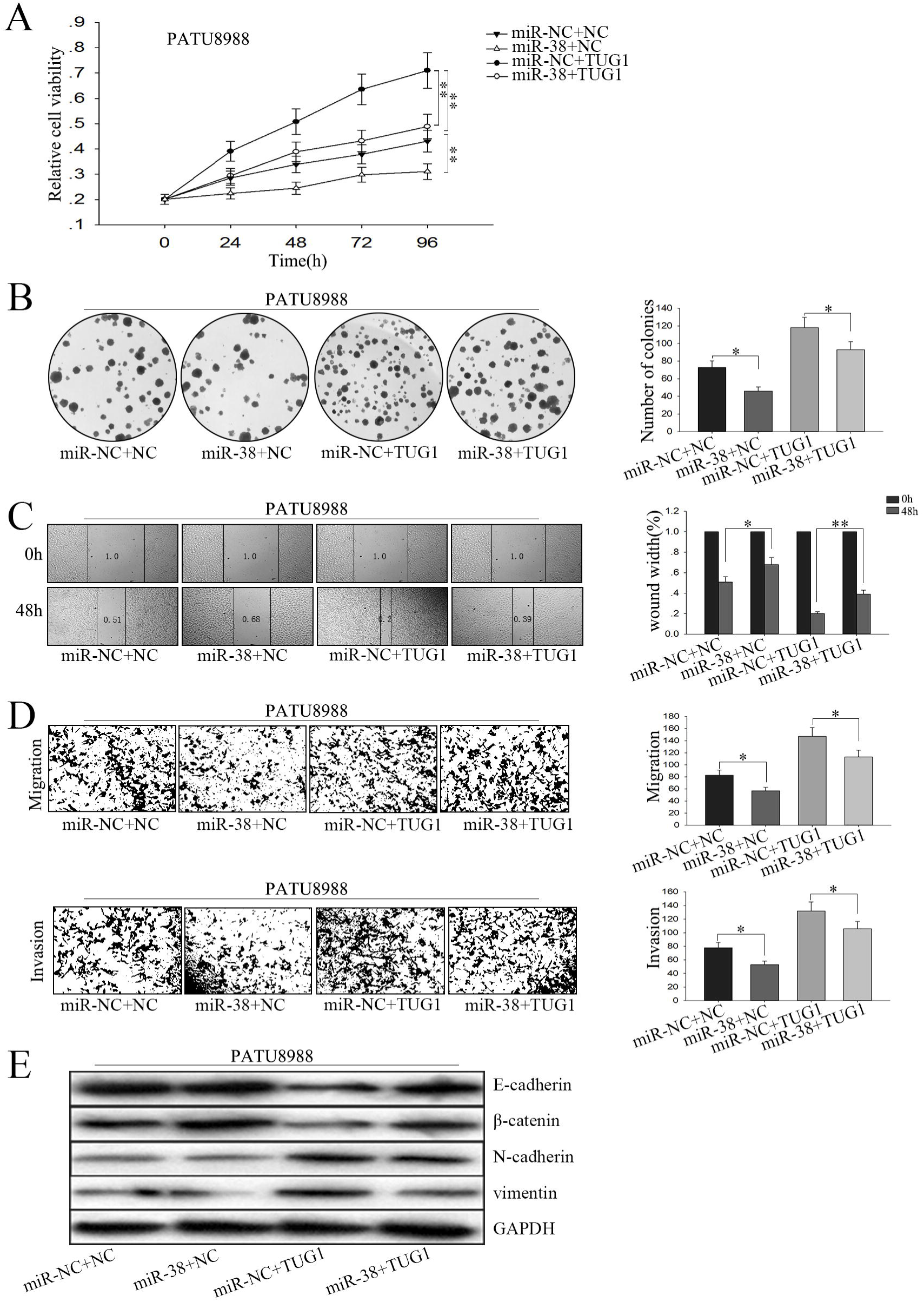
MiR-382 reversed the function of TUG1 in PC cells. **(A)** Cell MTT assay, **(B)** colony formation assay, **(C)** wound healing assay, **(D)** transwell assay **(E)** and western blot were performed in PATU 8988 cells transfected with miR-NC+NC, miR-NC+TUG1, NC+miR-382 and miR-382+ TUG1. All data were represented as the mean ± S.D. from three independent experiments, *P < 0.05, **P < 0.01.

### EZH2, a target of miR-382, is positively regulated by TUG1

Previously, we conducted gene expression array analysis using clinical samples resulting in the identification of several genes (Fig. 5A), which are involved in PC progression and metastasis. To screen the appropriate target of miR-382, we first measured the identified genes expression in the cells treated with miR-382 mimics. As present in Fig. 5B, miR-382 showed significant effect on the level of EZH2. And RT–qPCR showed that, in PC tissues, EZH2 was up-regulated in the tumor specimens, which was positively correlation with TUG1 (Fig. 5C). Additionally, bioinformatics analysis revealed that in the 3’UTR region of EZH2 contains a bind site for miR-382 (Fig. 5D left). Moreover, results from dual luciferase reporter assay showed that co-transfection of PANC-1 cells with pmirGLO-EZH2-WT vector and miR-382 mimic significantly reduced luciferase reporter activity, on the contrary, cells co-transfected with pmirGLO-EZH2-MUT vector and negative control showed a higher level of luciferase activity (Fig. 5D right). Furthermore, PATU 8988 and PANC-1 cells were transfected with miR-382 mimics and miR-382 inhibitor to decrease or increase miR-382 expression, respectively. Overexpression of miR-382 led to a significant down-regulation of EZH2, while knockdown of miR-382 significantly increased the level of EZH2 both in mRNA and protein levels (Fig. 5E). Additionally, results from RT-qPCR and western blot revealed that overexpression of TUG1 increased the level of EZH2 while down-regulated of TUG1 led to reduction of EZH2 both in miRNA and protein levels (Fig. 5F). Collectively, these findings suggest that EZH2 is a target of miR-382 and is positively regulated by TUG1.

**Figure 5.**
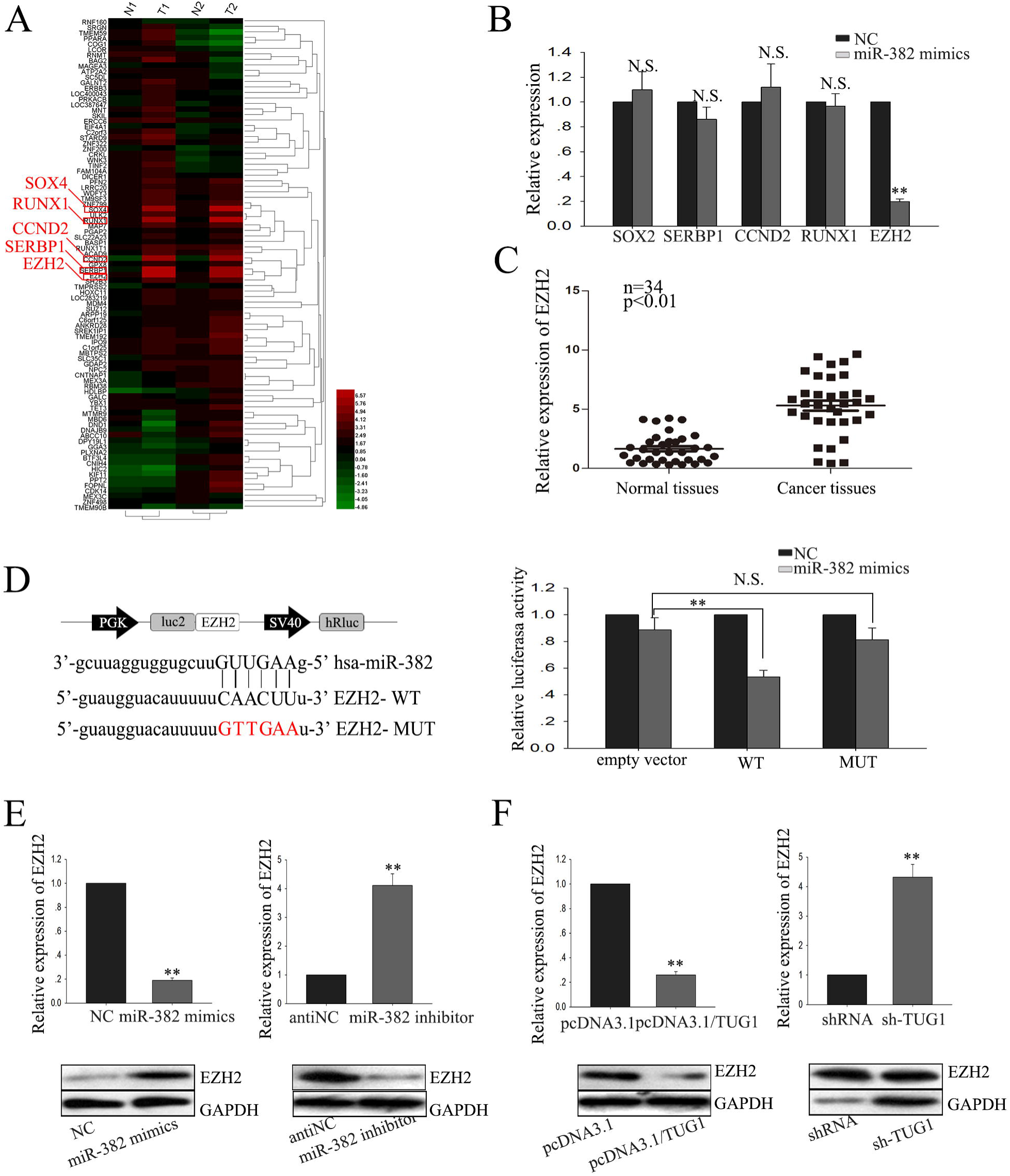
EZH2, a target of miR-382, is positively regulated by TUG1. **(A)** Gene expression array analysis of clinical samples. **(B)** MiR-382 showed significant effect on the level of EZH2. **(C)** EZH2 expression in pair samples of PC and adjacent normal tissues. **(D)** Luciferase assays of the indicated cells transfected with pmirGLO-EZH2-WT or pmirGLO-EZH2-MUT reporter and NC or miR-382 mimic. **(E)** The expression level of EZH2 both in mRNA and protein level in response to the level of TUG1and miR-382 **(F)** was measured. All data were represented as the mean ± S.D. from three independent experiments, *P < 0.05, **P < 0.01.

### Knockdown of EZH2 abolished the function of TUG1

We transfected PATU898 with TUG1expression vector and sh-EZH2 to study the effects of sh-EZH2 on cell migration and EMT mediated by TUG1. MTT and colony formation assay revealed that TUG1 promoted and sh-EZH2 suppressed cell proliferation, while co-transfection of sh-EZH2 and TUG1 expression vector showed that TUG1 promoted cell proliferation inhibited by sh-EZH2 (Fig. 6A-B). Would healing assay and transwell assay revealed that TUG1 promoted and sh-EZH2 suppressed cell migration, while co-transfection of sh-EZH2 and TUG1 expression vector showed that TUG1 promoted cell migration inhibited by sh-EZH2 (Fig. 6C-D). Moreover, results from western blot demonstrated that TUG1 promoted and sh-EZH2 suppressed PC cell EMT phenotype formation, while co-transfection of sh-EZH2 and TUG1 expression vector showed that sh-EZH2 reversed the EMT promoted by TUG1 to MET (Fig. 6E). Those findings suggest that downexpression of EZH2 inhibited PC cell migration and reversed EMT formation.

**Figure 6.**
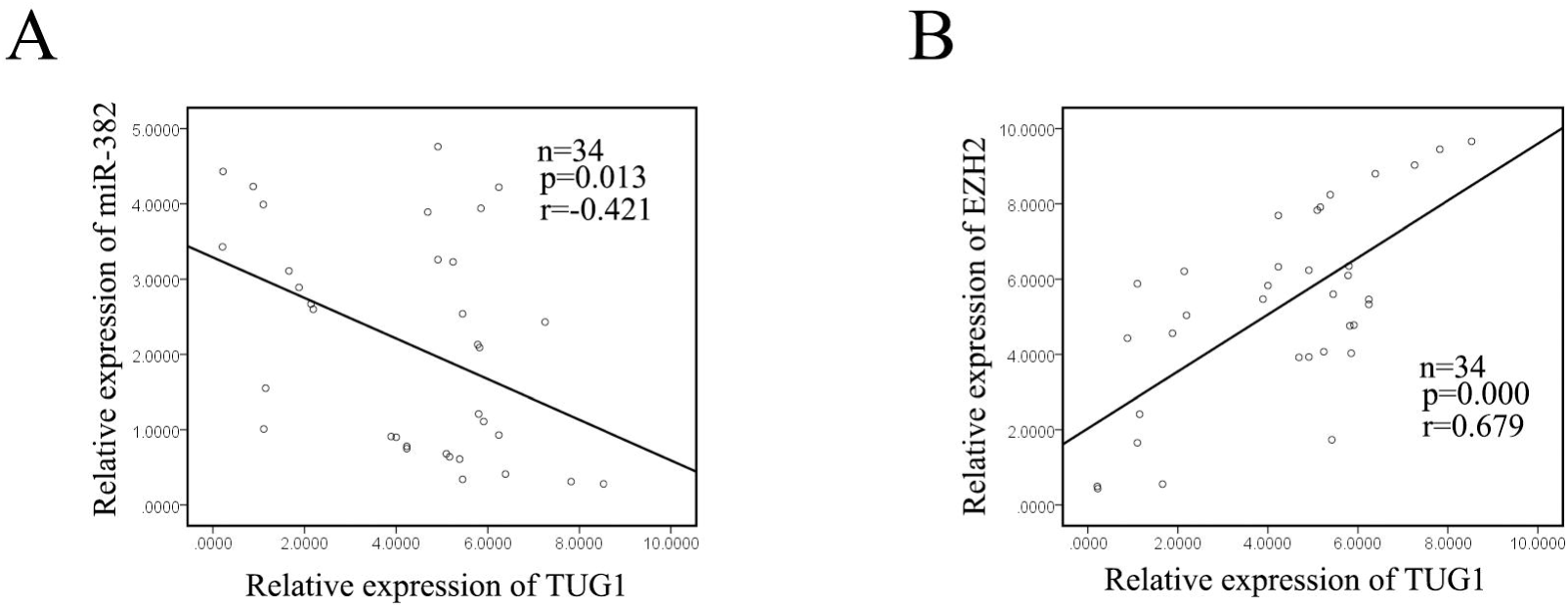
Knockdown of EZH2 abolished the function of TUG1. **(A)** Cell MTT assay, **(B)** colony formation assay, **(C)** wound healing assay, **(D)** transwell assay and **(E)** western blot were performed in PATU 8988 cells transfected with sh-RNA+NC, sh-EZH2+NC, sh-RNA+TUG1 and sh-EZH2+TUG1. All data were represented as the mean ± S.D. from three independent experiments, *P < 0.05, **P < 0.01.

### The expression level of TUG1 was negatively correlated with miR-382 and positively correlated with EZH2 in PC tissues

Finally, the relationship of the three molecules was analyzed. As shown in Fig. 7A, miR-382 levels and TUG1 levels were negatively correlated (2-tailed Spearman’s correlation, r=-0.421, p<0.01). What’s more, the level of EZH2 showed a positively correlated with TUG1 levels (2-tailed Spearman’s correlation, r=0.697, p<0.01; Fig. 7B). Collectively, these data indicated that there is a regulatory signaling pathway in which TUG1 regulates EZH2 by competitively sponging miR-382, inducing increased migration capacity and enhanced EMT phenotype formation PC cells.

**Figure 7.**
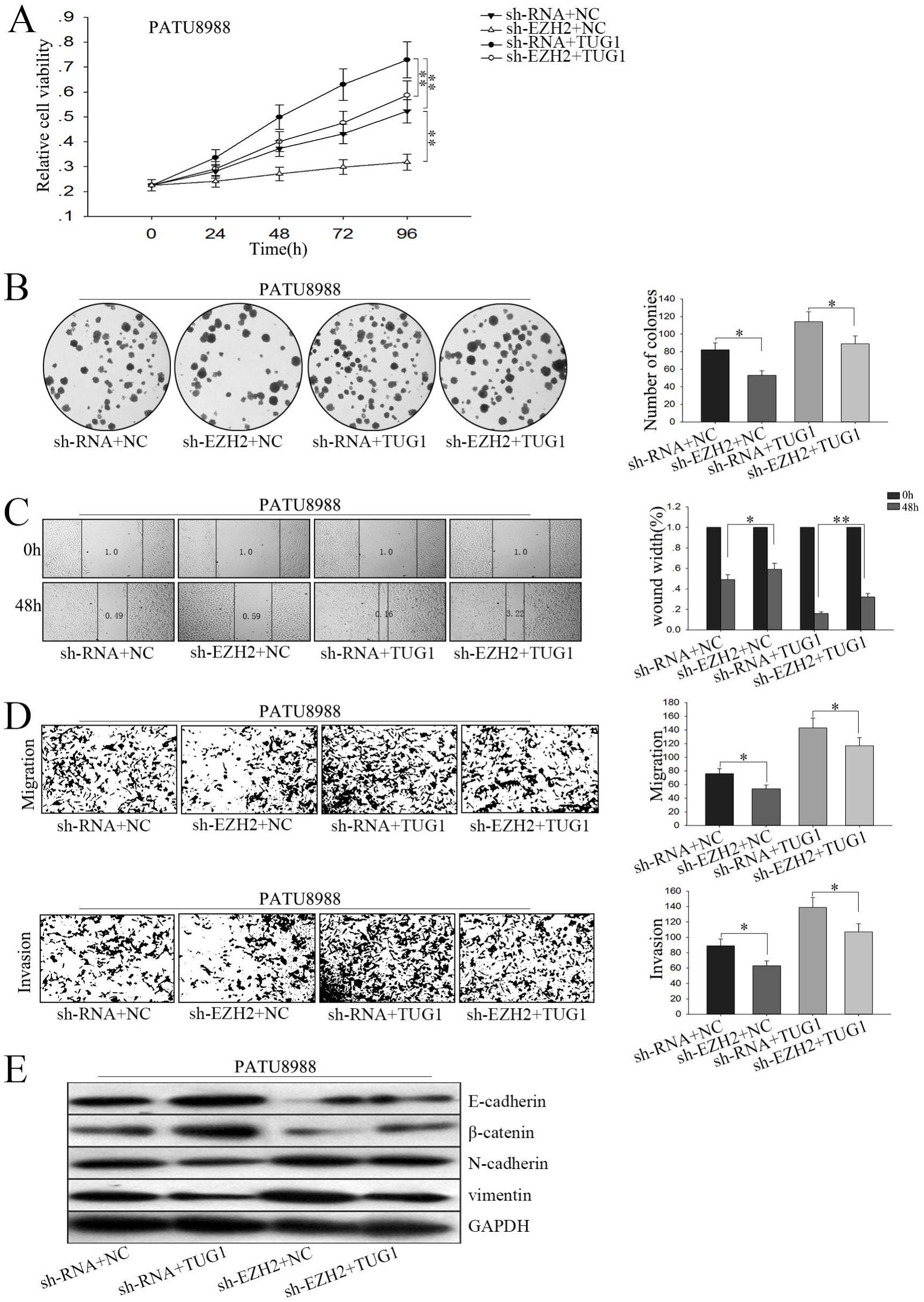
The expression level of TUG1 was negatively correlated with miR-382 and positively correlated with EZH2 in PC tissues. **(A)** miR-382 and **(B)** EZH2 expression levels were negatively (2-tailed Spearman’s correlation, r=-0.421, p<0.01) or positively (2-tailed Spearman’s correlation, r=0.679, p<0.01) correlated with TUG1 expression level in our expression cohort via qPCR.

## Discussion

LncRNAs have emerged as key regulators in cellular development and human diseases through modulating gene expression programs at transcriptional, post-transcriptional and epigenetic level; for instance, Koirala P et al. revealed that lncRNA AK023948 is a positive regulator of AKT; Xia M et al. demonstrated that lncRNA HOTAIR promotes metastasis of renal cell carcinoma by up-regulating histone H3K27 demethylase JMJD3; Xiong H et al. reported that lncRNA HULC triggers autophagy via stabilizing Sirt1 and attenuated the chemosensitivity of HCC cells[15-17]. Targeting lncRNAs-based signal pathway might be novel therapy methods. However, the roles of lncRNAs in PC carcinogenesis are not well understood. Investigating the underlying molecular mechanism by which lncRNAs function in PC might facilitate the exploitation of novel therapeutic targets. Despite many effort have been made in recent years, the survival rate of PC still remains unsatisfied. Therefore, investigating the mechanisms underlying the progression of PC might facilitate the development of novel treatments that improve patient prognosis.

LncRNA Taurine Up – regulated Gene 1 (TUG1) was initially identified as a transcript up - regulated by taurine, siRNA - based depletion of TUG1 suppresses mouse retinal development[18]. And the abnormal expression of TUG1 has been reported in many cancers[19-26]. In the current study, based on the results of lncRNA expression array analysis of clinical PC samples and RT–qPCR, we demonstrated that the lncRNA-TUG1 is upregulated in PC and serve as an oncogene in tumorigenesis.

Moreover, compared with the normal HPDE6-C cells, TUG1 expression was strikingly higher in four PC cell lines except PATU 8988 cell lines. Employing loss-of-function and gain-of-function approaches, we identified that TUG1 plays a key role in PC cell proliferation, migration and EMT formation. Deletion of TUG1 significantly weakened PC cell growth ability, decreased cell migration capacity and inhibited the EMT phenotype formation, and vice vasa in TUG1 forced expression cells. Our findings revealed that TUG1 may function as an oncogene in PC, and its overexpression facilitated PC development and progression. However, the detailed mechanism by which TUG1 functions in PC warrants further investigations.

It has been revealed that endogenous transcripts containing miRNA response elements (MREs) can co-regulate each other by acting as miRNA sponges or ceRNAs, forming large-scale regulatory networks across the transcriptome [27, 28]. Recently, accumulating lncRNAs have been demonstrated as endogenous decoy for specific miRNAs and regulate their function; for example, in 2017, Sun Y et al. demonstrated that deregulation of miR-183 promotes melanoma development via lncRNA MALAT1 regulation and ITGB1 signal activation; Yang S et al. in 2016, reported that construction of differential miRNA-lncRNA crosstalk networks based on ceRNA hypothesis uncover key roles of lncRNAs implicated in esophageal squamous cell carcinoma; and in 2016, Zhao L showed that lncRNA SNHG5/miR-32 axis regulates gastric cancer cell proliferation and migration by targeting KLF4[11, 29, 30]. In our study, we provide evidence that TUG1 may function as a ceRNA to competitively sponge miR-382 and influence the distribution on specific target. Consistent with the function of miR-382 mimic, deletion of TUG1 could suppress miR-382 target EZH2, whereas ectopically forced expressed TUG1 inhibits miR-382 function, leading to suppression of its target gene EZH2. Therefore, the influence of TUG1 on PC cells proliferation, migration and EMT formation could be attributed, at least partially, to function as a ceRNA competitively sponging miR-382.

General, we show that the TUG1 is upregulated in PC tissues and cell lines. Its effects on cell proliferation, migration and EMT formation indicated its oncogenic property in PC tumorigenesis. In our study, TUG1 acts as a molecular sponge for miR-382 and regulates its target EZH2. This reciprocal repression of TUG1 and miR-382 may highlight the important role of RNA–RNA interaction and provide novel insight into lncRNA-based mechanisms underlying various aspects of tumorigenesis.

## Conflicts of interest

No conflicts of interest to disclose.

## Acknowledgement

This study is supported by project co-sponsored by province and ministry (WKJ-ZJ-1706) and the technology innovation team of diagnosis and treatment of abdominal surgery of Wenzhou, Zhejiang Province, China (2016-354) and Zhejiang Provincial Natural Science Foundation of China (No. LY15H160056).

